# An Efficient Approach for SARS-CoV-2 Monoclonal Antibody Production via Modified mRNA-LNP Immunization

**DOI:** 10.1101/2022.04.20.488878

**Authors:** Fu-Fei Hsu, Kang-Hao Liang, Monika Kumari, Wan-Yu Chen, Hsiu-Ting Lin, Chao-Min Cheng, Mi-Hua Tao, Han-Chung Wu

## Abstract

Throughout the COVID-19 pandemic, many prophylactic and therapeutic drugs have been evaluated and introduced. Among these treatments, monoclonal antibodies (mAbs) that bind to and neutralize SARS-CoV-2 virus have been applied as complementary and alternative treatments to vaccines. Although different methodologies have been utilized to produce mAbs, traditional hybridoma fusion technology is still commonly used for this purpose due to its unmatched performance record. In this study, we coupled the hybridoma fusion strategy with mRNA-lipid nanoparticle (LNP) immunization. This time-saving approach can circumvent biological and technical hurdles, such as difficult to express membrane proteins, antigen instability, and the lack of posttranslational modifications on recombinant antigens. We used mRNA-LNP immunization and hybridoma fusion technology to generate mAbs against the receptor binding domain (RBD) of SARS-CoV-2 spike (S) protein. Compared with traditional protein-based immunization approaches, inoculation of mice with RBD mRNA-LNP induced higher titers of serum antibodies. In addition, the mAbs we obtained can bind to SARS-CoV-2 RBDs from several variants. Notably, RBD-mAb-3 displayed particularly high binding affinities and neutralizing potencies against both Alpha and Delta variants. In addition to introducing specific mAbs against SARS-CoV-2, our data generally demonstrate that mRNA-LNP immunization may be useful to quickly generate highly functional mAbs against emerging infectious diseases.

## INTRODUCTION

As the world continues to grapple with the coronavirus COVID-19 pandemic (Zhu N, et al., 2020; Sun P, et al., 2020), monoclonal antibodies (mAbs) have become increasingly used in applications such as basic research, diagnosis, therapeutics for SARS-CoV-2 infection (Hwang YC, et al., 2022; Chapman AP, et al., 2021). Fortunately, the hybridoma technology developed by Georges Kohler and Cesar Milstein in 1975 has made it possible to obtain large numbers of mAbs for these purposes (Köhler G, et al., 1975). Despite its widespread application, hybridoma technology still suffers from some limitations. For example, the conventional methodology used for hybridoma generation involves multiple injections of a protein antigen with or without adjuvant (Chiarella P, et al., 2008). It is sometimes a major challenge to prepare high-quality protein antigen for immunization, yet this step is necessary to generate high-precision mAbs that are able to recognize the native viral antigen (Takeda H, et al., 2015). Purification of protein antigens is also a time consuming and labor-intensive process, as optimized protocols must be created for each target (Holzlöhner P, et al., 2017). Moreover, it is technically difficult to express or purify certain antigens, such as transmembrane, glycosylated, toxic, or unstable proteins (Parray HA, et al., 2020; Zuo X, et al., 2005). Once the antigen is produced, the purified recombinant proteins are often mixed into adjuvant formulations that might alter the native protein conformations and lead to unexpected immune responses and undesired consequences in the immunized animals (Parray HA, et al., 2020). In addition, it is well known that maintenance of structural integrity of immunogens is a critical concern for the induction of functional mAbs (Liu S, et al., 2016). For all of these reasons, the use of protein antigens is often inadequate, and an alternative approach of expressing intact immunogens *in vivo* may be better suited for inducing mAbs with desired biological activities.

In recent years, the ability to produce mRNA in a cell-free environment by *in vitro* transcription (IVT) has provided a realistic opportunity to generate any protein of interest in cells or animals (Maruggi G, et al., 2019). However, the use of mRNA to induce protein production in animals is complicated by its susceptibility to degradation by nucleases, inherent instability, stimulation of excessive inflammatory responses and inefficient *in vivo* delivery (Hou X, et al., 2021; Wang Y, et al., 2021). During the last two decades, intensive research has been carried out to develop effective strategies for stabilization and delivery mRNA (Jackson NAC, et al., 2020; Pardi N, et al., 2018). As a result of these efforts, co-formulation into lipid nanoparticles (LNPs) has become one of the most successful and commonly used methods for mRNA delivery *in vivo* (Ickenstein LM, et al., 2019). Because of its reliability, mRNA-LNPs are being extensively utilized in a broad range of new and potential treatments, such as regenerative medicines, vaccines, immunotherapies, and gene editing applications (Baptista B, et al., 2021).

In this study, we designed and developed an LNP-encapsulated nucleoside-modified SARS-CoV-2 RBD mRNA (amino acids [aa] 319–541) with Ig kappa chain leader sequence for protein secretion (called RBD mRNA-LNP). The expression levels and secretory efficiencies of RBD mRNA-LNP were initially evaluated in HEK293T cells by Western blotting and ELISA. We then immunized mice with RBD mRNA-LNP over a period of 7 weeks and found that the animals had higher serum antibody titers than mice vaccinated with RBD protein in adjuvant over a period of 10 weeks. After using the mice for hybridoma production, we identified six different IgG mAbs, demonstrating the applicability of this novel immunization approach for mAb generation. Each of the six mAbs was evaluated in terms of its target binding affinity and neutralization potential using ELISA and pseudovirus neutralization assays. The mAbs were all found to exhibit specific binding to SARS-CoV-2 RBD protein and at least some of the tested variants. Among the six mAbs, RBD-mAb-3 showed remarkably potent neutralization against both Alpha and Delta variants (Alpha, IC_50_ = 10.99 ± 1.45 ng/ml and Delta, IC_50_ = 11.12 ± 1.09 ng/ml). Binding of RBD-mAb-3 to S protein RBD was inhibited by mutations at sites Y453A and Q474A. Together, these data suggest that mRNA-LNP immunization has the potential to overcome or bypass many current challenges in mAbs development, allowing fast turnaround times that can benefit efforts to monitor or treat emerging infectious diseases.

## RESULTS

### Design and characterization of the RBD mRNA-LNP immunogen for SARS-CoV-2

Lipid nanoparticles (LNPs) are one of the most commonly used tools for *in vivo* delivery of mRNA (Ickenstein LM, et al., 2019). To develop an mRNA-LNP to deliver the RBD of SARS-CoV-2 as an immunogen, the RBD of SARS-CoV-2 (amino acids [aa] 319–541) was produced by IVT from a template based on the published sequence of the virus (Figure 1A). This synthetic mRNA harbored an optimal CAP-1 structure and a signal peptide sequence from human Ig κappa chain (Igκ). N1-methyl-pseudouridine (m1Ψ) was substituted for uridine throughout the mRNA sequence to increase mRNA stability and translation (Parr CJC, et al., 2020). The LNP was composed of four types of lipids, including D-Lin-MC3-DMA, DSPC, cholesterol and PEG-lipid at molar ratios of 50:10:38.5:1.5 (Figure 1B). After production, the RBD mRNA-LNP samples were observed by cryo-electron microscopy (Figure 1C), which revealed the formulation was comprised of spherical and homogenously distributed LNPs without aggregation. As shown in Figure 1D, agarose gel retardation was performed in presence and absence of Triton X-100. mRNA encapsulated in the LNP particles (lane 2) showed no free mRNA band, suggesting that the encapsulation of mRNA into the LNPs was complete. Following the addition of Triton X-100 to disrupt the lipid membrane (lane 3), the mRNA was released from the LNPs but did not exhibit any degradation. Since the mRNA released from the disrupted LNP was at the same molecular weight as the naked mRNA, we concluded that the integrity of mRNA was not altered by encapsulation. The mean particle size distribution of RBD-mRNA-LNP ranged from 70 to 80 nm and PDI was less than 0.15, indicating a mono-dispersed size distribution (Figure 1E). Of note, the positive surface charge of LNPs may allow for their interaction with negatively charged plasma membrane to facilitate cellular uptake. In our formulation, the particle size distribution and surface charge were suitable for cellular uptake by endocytosis pathways.

**Figure 1.**
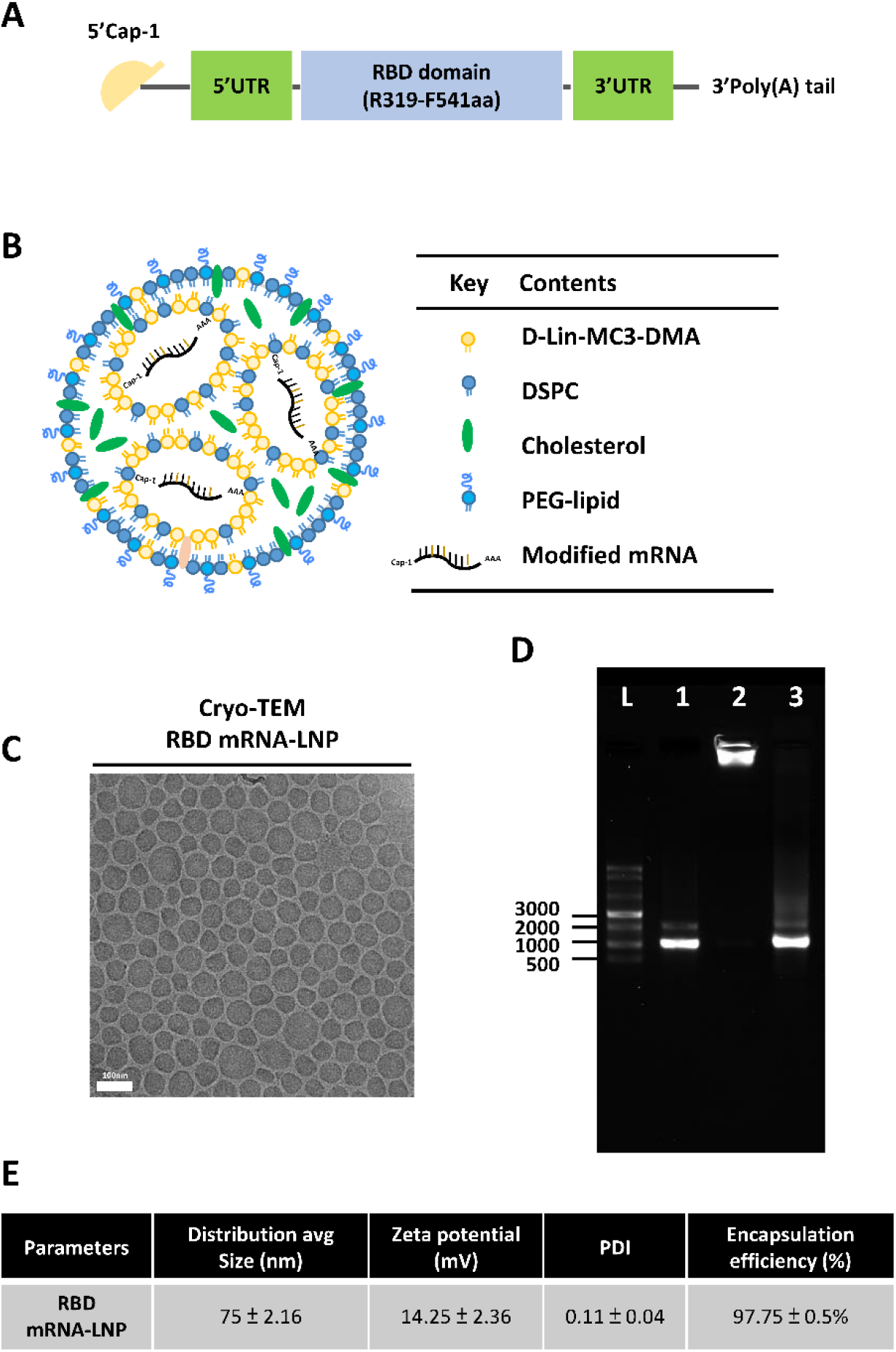
Design and physicochemical characterization of the modified RBD mRNA-LNP used for immunization. (A) Schematic illustration of SARS-CoV-2 RBD mRNA; comprises a 5’ cap, 5’ UTR, signal peptide, antigen (RBD of the spike protein), 3’ UTR, and 3’ poly(A) tail. (B) Schematic representation of the LNP-encapsulated RBD mRNA. (C) Cryo-TEM imaging was performed on RBD mRNA-loaded LNPs; the LNPs consisted of D-Lin-MC3-DMA:DSPC:Cholesterol:DMG-PEG2000 at 50:10:38.5:1.5 molar ratios. Scale bar, 100 nm. (D) Agarose gel electrophoresis of RBD mRNA-LNP was performed with and without 1% Triton X-100 to assess mRNA integrity. Naked mRNA was used as a control. L = single-stranded RNA ladder, 1 = RBD mRNA alone, 2 = RBD mRNA-LNP (in 1X TE), 3 = disrupted RBD mRNA-LNP (in 1% Trition X-100). (E) Average size, zeta potential and PDI distribution were measured with dynamic light scattering (DLS); Encapsulation efficiency (EE%) was determined with a Ribogreen assay kit.

### *In vitro* characterization of RBD mRNA or RBD mRNA-LNP expression

To assess cell entry, cargo escape and translation of the transported mRNA molecules, RBD protein expression was measured after transfecting HEK293T cells with RBD mRNA-LNP. Two days after transfection, RBD protein secreted into the medium was analyzed by Western blotting (Figure 2A) and ELISA (Figure 2B). Negative controls included media from non-transfected cells. The results demonstrated that the RBD could be efficiently produced and secreted by HEK293T cells into the culture supernatant. In addition, we evaluated the translation efficiency of RBD mRNA after lipofectamine-mediated transfection into cell cultures (Supplementary Figure 1).

**Figure 2.**
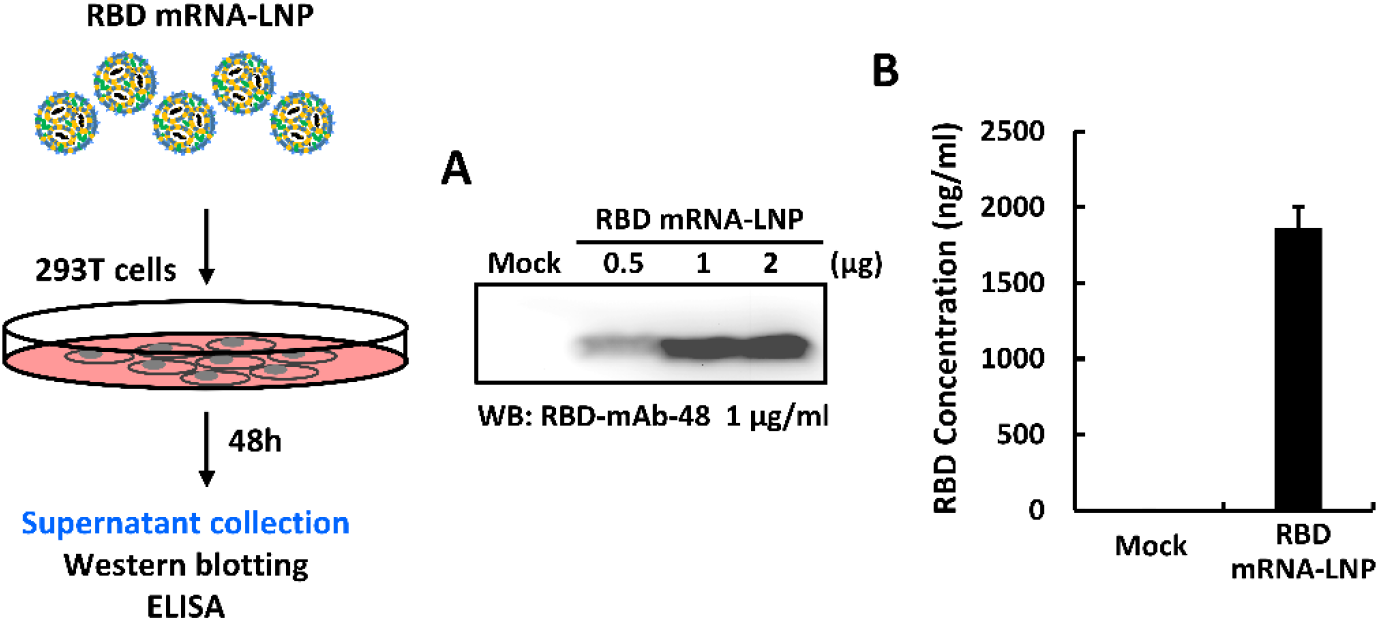
*In vitro* characterization of immunogen expressed from RBD mRNA-LNP. Schematic of the RBD mRNA-LNP transduction protocol for 293T cells. (A, B) RBD protein expression level was measured in HEK293T cells transfected with RBD mRNA-LNP by Western blotting (A) and ELISA (B), using an anti-RBD antibody. Mock transfected HEK293T cells served as a negative control.

### Immunization with RBD mRNA-LNP elicits a robust antigen-specific immune response in mice

Next, we wanted to compare the strengths and durations of antibody responses between mice immunized with RBD mRNA-LNP and those subjected to a traditional protein-based approach. BALB/c mice were immunized via intramuscular injection with RBD mRNA-LNP at 2-week intervals or received intraperitoneal injection with recombinant RBD protein in CFA-IFA adjuvant at 3-week intervals. A schematic representation of the immunization schedule is shown in Figure 3A. Pre-immune and immune serum were respectively collected before the priming injection (week 0) and one week after the second booster. Subsequently, the RBD-binding antibody titers of the serum samples were evaluated by ELISA. Sera from mice immunized with the RBD mRNA-LNP exhibited significantly higher titers against RBD protein, as compared to the mice immunized with the protein and adjuvant (Figure 3B). This result suggests that RBD mRNA LNP is highly immunogenic in mice and is suitable for subsequent production of mAbs against RBD. As such, the RBD mRNA-LNP-immunized mice were used for cell fusion. One week after the final injection, splenocytes were harvested from the mice and fused with NS1/1-Ab4-1 (NS-1) mouse myeloma cells. The resulting fused hybrid cells were cultured and selected in HAT tissue culture media.

**Figure 3.**
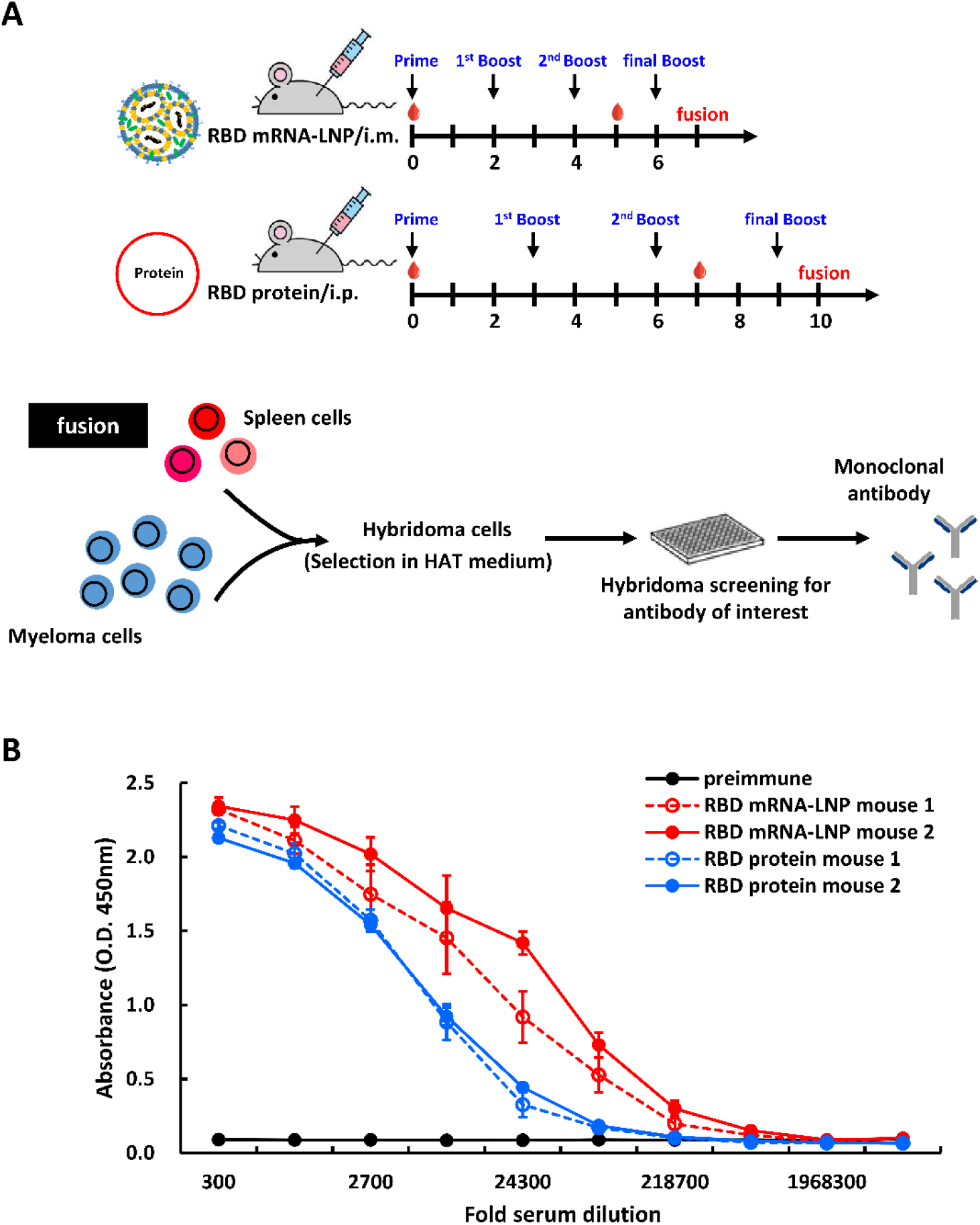
Immunization with RBD mRNA-LNP elicits a robust antigen-specific immune response in mice. (A) Schematic of immunization regimens in mice. Female BALB/c mice were immunized by three i.m. injections of RBD mRNA-LNP at 2-week intervals or by three i.p. injections of recombinant RBD protein at 3-week intervals. Red droplets indicate blood draws. After the final booster, the spleens were harvested and fused with NS1 cells to generate hybridomas, which were screened for RBD-mAbs. (B) ELISA was used to determine the titers of preimmune and hyperimmune sera against RBD mRNA-LNP or RBD protein after three immunizations. Mouse sera were collected before immunization and after the third immunization and incubated with RBD protein-coated plates. The absorbance values were read at O.D. 450nm.

### Generation and characterization of monoclonal antibodies

Supernatants of hybridoma clones were screened by ELISA for binding to recombinant RBD protein. As shown in Figure 4A, six hybridoma culture supernatants containing anti-RBD IgG1 (κ light chain) showed specific dose-dependent binding to RBD protein. Nevertheless, like many other RNA viruses, SARS-CoV-2 has a high mutation rate, and some mutations may cause new variants to be more transmissible and virulent than the original SARS-CoV-2 wild-type strain, reported in late 2019 (Harvey WT, et al., 2021). Over the course of the pandemic, five SARS-CoV-2 lineages have been designated by the World Health Organization as variants of concern (VOCs), including Alpha (B.1.1.7), Beta (B.1.351), Gamma (P1) and Delta (B.1.617.2) and the recent Omicron variant (B.1.1.529). Therefore, we also tested the binding capabilities of our hybridoma culture supernatants to RBDs with mutations corresponding to those found in the five VOCs (Figure 4B and Supplementary Figure 2A). RBD-mAb-1 and RBD-mAb-5 recognized wild-type RBD and RBD variants with mostly similar binding activities, but showed relatively weaker binding to the Beta RBD variant. Notably, RBD-mAb-1 showed efficient binding to the Omicron RBD variant, while RBD-mAb-5 had much weaker binding activity for the Omicron RBD variant. RBD-mAb-2, RBD-mAb-4 and RBD-mAb-6 bound well to wild-type RBD and most RBD variants, but their binding to Omicron RBD was relatively weak. RBD-mAb-3 bound strongly to wild-type RBD, Alpha RBD, and Delta RBD, but had no detectable binding to the Beta, Gamma and Omicron variants. In addition to evaluating the binding of hybridoma supernatants to recombinant proteins, we examined the neutralizing potencies of our six hybridoma culture supernatants toward pseudoviruses of the five SARS-CoV-2 VOCs. Consistent with the ELISA data showing that RBD-mAb-3 bound to Alpha and Delta RBD variants, the pseudovirus neutralization assays revealed that RBD-mAb-3 exhibited high neutralizing abilities for Alpha and Delta variants, with IC_50_ values of less than 6.43 ng/ml (Figure 4C). While this selected hybridoma clone was effective against the virulent Delta strain, it did not effectively neutralize the Omicron pseudovirus (Supplementary Figure 2B).

**Figure 4.**
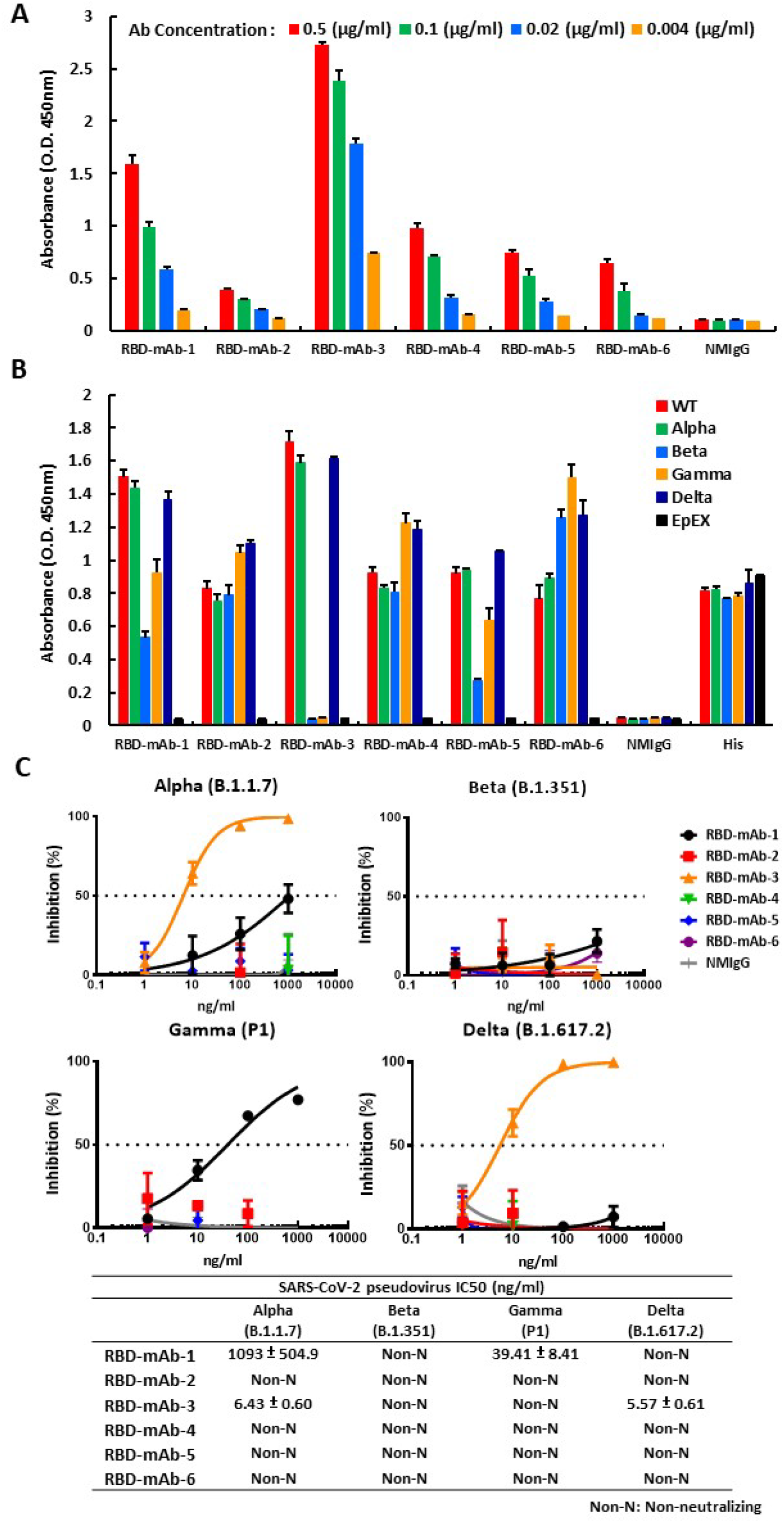
Generation and characterization of mAbs from hybridomas. (A) Hybridomas were generated, and serial dilutions of the cell culture supernatants were screened for RBD WT-His binding ability by ELISA. (B) The binding affinities of hybridoma culture supernatants were also assessed for RBDs of different SARS-CoV-2 variants by ELISA. EpEX protein served as a negative control. (C) Neutralization assays were performed with hybridoma culture supernatants targeting four SARS-CoV-2 VOC pseudoviruses. Assays were performed in triplicate; each point represents the mean ± SEM. IC_50_ values were calculated with GraphPad Prism software.

### Neutralizing effect of RBD-mAb-3 against SARS-CoV-2 Alpha and Delta

As RBD-mAb-3 showed efficient neutralization of Alpha and Delta SARS-CoV-2 variant pseudoviruses, it was selected for further IgG antibody purification and validation. To detect the binding activities of purified RBD-mAb-3, ELISA was carried out. As shown in Figure 5A, RBD-mAb-3 had high binding potencies against RBD Alpha and Delta variants. The K417, Y453, Q474, F486, Q498, T500, and N501 residues within the RBD are known to be responsible for S protein contact with the host receptor, ACE2 (Yan R, et al., 2020). A cellular ELISA-based binding assay was performed to evaluate whether RBD-mAb-3 binding is affected by alanine mutations at positions K417, Y453, Q474, F486, Q498, T500 and N501 (Figure 5B). The results showed that Y453A and Q474A mutations significantly decreased the binding of RBD-mAb-3, suggesting that these residues contribute to the RBD-mAb-3 epitope. We then further examined the neutralizing abilities of purified RBD-mAb-3 toward SARS-CoV-2 Alpha and Delta pseudoviruses. Notably, RBD-mAb-3 exhibited high neutralizing capacities for Alpha and Delta variants, with respective IC_50_ values of 10.99 and 11.12 ng/ml (Figure 5C). Given the specificity and binding affinity of RBD-mAb-3 towards the Delta RBD variant, we also tested its neutralization potential in a plaque reduction neutralization test (PRNT) using authentic SARS-CoV-2 Delta variant (Figure 5D). RBD-mAb-3 effectively neutralized viral infection with a PRNT_50_ value of 12.71 ng/ml. Collectively, these data indicate that RBD-mAb-3 is a potent neutralizing mAb for SARS-CoV-2 Delta variant.

**Figure 5.**
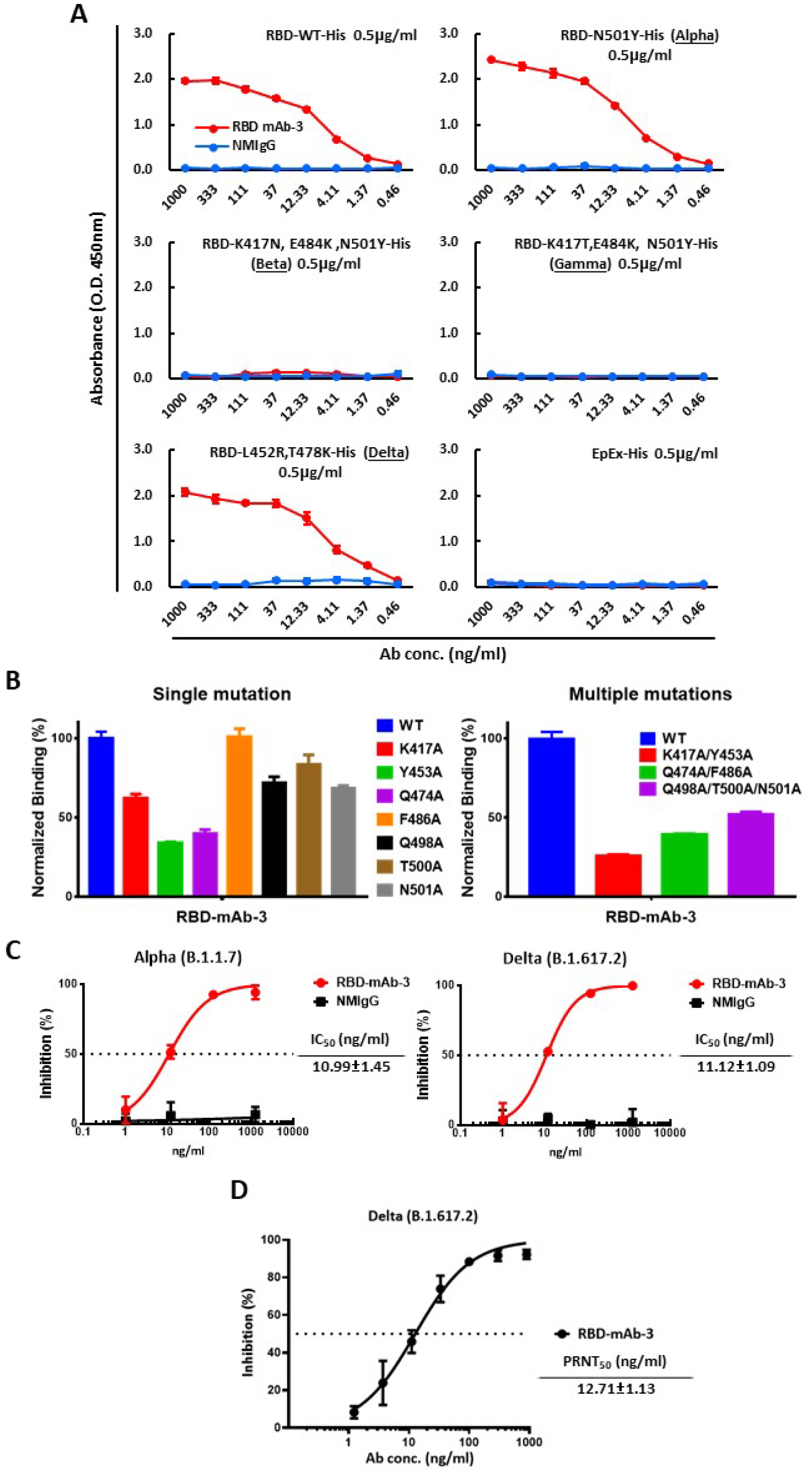
RBD-mAb-3 demonstrates potent binding and neutralization of SARS-CoV-2 Delta. (A) The binding of RBD-mAb-3 to different SARS-CoV-2 VOC RBDs was determined by ELISA analysis. EpEX served as a negative control. (B) Epitope mapping of RBD-mAb-3 using mutagenesis. RBD-mAb-3 binding to mutant RBDs with single or multiple alanine mutations was normalized to its binding to wild-type (WT) RBD; measurements were made with cellular ELISA. (C) Neutralization assays were performed on SARS-CoV-2 Alpha and Delta pseudoviruses with purified RBD-mAb-3. Assays were performed in triplicate; each point represents the mean ± SEM. IC_50_ values were calculated with GraphPad Prism software. (D) RBD-mAb-3 neutralizes SARS-CoV-2 Delta according to PRNT. The PRNT_50_ value was calculated with GraphPad Prism software. Assays were performed in triplicate, and points represent the mean ± SD.

## DISCUSSION

Throughout the SARS-CoV-2 pandemic, mAbs have gained widespread use in treatments and diagnostic tools. For example, several neutralizing mAbs have been granted emergency use authorizations (EUAs) for treatment of non-hospitalized patients with mild-to-moderate COVID-19 (Hwang YC, et al., 2022; Taylor PC, et al., 2021). Since hybridoma technology is one of the most reliable approaches for mAb production, it has been widely applied in many contexts (Lu RM, et al., 2020). However, preparation of traditional protein antigens is often the most critical and time-consuming step in the mAb production process, especially for mAbs against hard-to-express and -purify antigens like membrane proteins, glycoproteins or conformational epitopes. Therefore, mRNA-LNP immunization may be a more effective strategy to quickly produce mAbs against such targets. The COVID-19 pandemic has accelerated the research toward mRNA technologies (Kumar A, et al., 2022), and the success of mRNA-LNP vaccines suggests that the technology may be efficiently applied for the rapid development of high-quality mAbs against SARS-CoV-2 and other pathogens. In this study, we demonstrated this potential by using RBD mRNA-LNP as a novel immunogen for hybridoma-based mAb production.

The main advantage of mRNA-LNP immunization is that the mRNA is easily produced by IVT and amenable to rapid manipulation of antigens. The only custom-produced material necessary for this method is the plasmid encoding an optimized target sequence. In addition, our LNPs include D-Lin-MC3-DMA, a commercially available cationic lipid, which can facilitate efficient encapsulation and delivery of RBD mRNA to cells *in vitro* and *in vivo*. Most importantly, even after antigen is produced, the process of generating hybridoma-based mAbs normally takes an average of 6 to 8 months to complete, with animal immunization being the most time-consuming step. Using the mRNA-LNP immunization strategy, we were able to successfully generate several hybridomas secreting antigen-specific mAbs while reducing the time from initial immunization to spleen harvest from 10 to 7 weeks. In the future, application of mRNA-LNP immunization to human antibody-producing transgenic mice (Murphy AJ, et al. 2014; Lu RM, et al., 2020) coupled with high throughput screening of single B cells may be an even more time-efficient strategy for generating of mAbs to be used in therapeutics or diagnostics.

## MATERIALS AND METHODS

### Production of modified IVT mRNA

The DNA template contained a T7 promoter followed by a codon optimized wild-type (Wuhan-Hu-1, GenBank YP_009724390.1) RBD sequence of SARS-CoV-2 S protein (Arg319-Phe541) flanked by the 5’ UTR, IgG kappa leader sequence and the 3’ UTR of the α-globin gene. A unique EcoRV site was included downstream of the 3’ UTR poly(A) tail region. Prior to the IVT reaction, the plasmid was linearized using the EcoRV and purified with the NucleoSpin Gel and PCR Clean-up Kit (Macherey & Nagel Co. Düren, Germany). mRNA was synthesized by IVT according to the manufacturer’s recommendations, using HiScribe T7 (NEB, MA, USA) with co-transcriptional CleanCap ®AG (Trilink, CA, USA) and N1-methyl-pseudouridine (Trilink, CA, USA). Synthesized mRNA was purified by DNase I (NEB, MA, USA) digestion followed by LiCl (Invitrogen, Thermo Fisher Scientific, Waltham, MA, USA) precipitation and 70% ethanol wash. To remove dsRNA from the transcribed mRNA, selective adsorption of the dsRNA to cellulose (Sigma-Aldrich, MA, USA) was performed in an ethanol-containing buffer. Purified mRNA was kept at -80 °C until further use

### Preparation of RBD mRNA-LNPs

Lipid-nanoparticle (LNP) formulations were prepared using a previously described method (Lee IJ, et. al., 2022). Briefly, lipids were solubilized in ethanol containing cationic lipid (DLin-MC3-DMA) (MedChemExpress, NJ, USA), 1, 2-distearoyl-sn-glycero-3-phosphocholine (DSPC) (Avanti Polar Lipids, NY, USA), cholesterol (Sigma, MA, USA) and PEG-2000 (MedChemExpress, NJ, USA) at molar ratios of 50:10:38.5:1.5. The lipid mixture was combined with 25 mM citrate buffer (pH 4.5) containing mRNA at a ratio of 1:3 using NanoAssemblr® IGNITE NxGen Cartridges (Precision NanoSystems Inc., BC, Canada). LNP-encapsulated mRNA samples were dialyzed against PBS (pH 7.4) and then concentrated using Amicon Ultra Centrifugal Filters (10 K MWCO; Millipore, Burlington, MA, USA) before being passed through 0.45-μm filters.

### Characterization of RBD mRNA-LNPs

The particle size distribution, polydispersity index (PDI) value and zeta potential of RBD mRNA-LNPs were analyzed by Dynamic Light Scattering (DLS, Zetasizer Nano ZS, Malvern Panalytical Ltd., Malvern, WR, UK). The sample was diluted 100-fold and equilibrated for 120 s at 25°C prior to size and zeta potential measurements. The hydrodynamic diameter (z-average) and zeta potential of RBD mRNA-LNPs were analyzed by Zetasizer software, version 7.11 (www.malvern.com). The morphology of RBD mRNA-LNPs in a dry state was observed using cryogenic transmission electron microscopy (cryo-TEM, Tecnai F20, Philips, Eindhoven, the Netherlands). Briefly, the sample solution was diluted 10-fold and transferred onto 300-mesh copper grids covered with porous carbon film (HC300-Cu, PELCO) for blotting and plunging in a 100% humidity temperature-controlled chamber using a Vitroblot system (FEI). The copper grids were stored in liquid nitrogen and transferred to the electron microscope on a cryo-stage for imaging. The mRNA encapsulation efficiency (EE%) and concentration was determined with a Quant-iT RiboGreen RNA assay kit (Invitroge, Thermo Fisher Scientific, Waltham, MA, USA). The agarose gel retardation assay was performed to analyze the size and integrity of bound/unbound mRNA. The bound/unbound mRNA in LNP nanoparticles was analyzed in the presence and absence of 1% Triton X-100.

### In vitro RBD expression of RBD mRNA and mRNA-LNPs

RBD mRNA was transfected into 293T cells via Lipofectamine™ 2000 Transfection Reagent (Invitrogen, Thermo Fisher Scientific, Waltham, MA, USA), while RBD mRNA-LNP was added directly to cells cultured in DMEM containing 10% FBS. The supernatants were collected at 48 h after transfection, and RBD expression was monitored by Western blot and sandwich ELISA. Sandwich ELISA was used to quantitatively measure the amount of RBD protein in cell culture media. Briefly, a mouse anti-RBD mAb was diluted to a concentration of 2 μg/ml in 0.1 M sodium carbonate buffer (NaHCO_3_/Na_2_CO_3_; pH 9.5). Then, a 96-well plate was coated with coating buffer at 4°C overnight. The next day, the coating buffer was discarded, and the plate was blocked with PBS containing 1% bovine serum albumin (BSA) at room temperature for 1 h. The coated plates were washed three times with PBS and incubated with calibrator solutions or serial dilutions of cell culture media at RT for 1 h. After washing the plates three times with 0.1% Tween-20 in PBS (PBST_0.1_), anti-RBD chAb diluted in 1% BSA/PBS at 1:2000 was applied at room temperature for 1 h. The plates were washed with PBST_0.1_ three times, then incubated with horseradish peroxidase (HRP)-conjugated anti-human IgG (Sigma, MA, USA) secondary antibody at room temperature for 1 h. After three washes with PBST_0.1_, signal was generated using 3,3’5,5’-Tetramethylbenzidine (TMB) substrate (TMBW-1000-01, SURMODICS). The reaction was stopped with 3 N HCl, and absorbance was measured at 450 nm by ELISA reader (Versa Max Tunable Microplate Reader; Molecular Devices).

### Mouse immunization

All procedures involving animals were performed in accordance with guidelines set by the Institutional Animal Care and Use Committee (IACUC) at Academia Sinica, Taiwan. BALB/c mice 6 to 8 weeks of age were immunized by four intramuscular injections of 10 μg RBD mRNA-LNP at 2-week intervals or four intraperitoneal injections of 50 μg recombinant RBD protein in CFA-IFA adjuvant at 3-week intervals. Serum samples were collected from each mouse before the priming injection and at one week after the final immunization.

### Generation of SARS-CoV-2 RBD-specific murine mAbs

Hybridomas secreting anti-RBD mAbs were generated and identified according to standard procedures (Köhler G, et al., 1975) with some modifications (Chen YC, et al., 2007; Tung KH, et al., 2013). Briefly, the spleen of an immunized mouse was removed, and splenocytes were fused with NSI/1-Ab4–1 (NS-1) myeloma cells. The fused cells were cultured in DMEM supplemented with 15% FBS, hypoxanthine-aminopterin-thymidine (HAT) medium and hybridoma cloning factor, in 96-well tissue culture plates. Hybridoma colonies were screened by ELISA for secretion of mAbs that bound to RBD recombinant protein. Selected clones were subcloned by limiting dilutions. Final hybridoma clones were isotyped using an isotyping kit. Ascitic fluids were produced in IFA-primed BALB/c mice. Hybridoma cell lines were grown in DMEM with 10% heat-inactivated FBS. The candidate mAbs were affinity purified with protein G sepharose 4B gels. ELISA assays were used to measure the activity and specificity of each antibody.

### Antibody binding to SARS-CoV-2 RBD protein variants by ELISA

Recombinant His-tagged RBD proteins from SARS-CoV-2 variants were purchased as lyophilized powders from ACROBiosystems Inc. (Newark, DE, USA). The 96-well ELISA plates were coated with 0.5 μg/ml SARS-CoV-2 variant RBD-His or EpEX-His (negative control) proteins in 0.1 M sodium carbonate buffer at 4°C overnight, followed by blocking with 1% BSA in PBS at room temperature for 2 h. After blocking, the coated plates were washed twice with PBS and stored at -20°C until further use. The protein contents of the culture supernatants from hybridomas or antibodies were quantified by the BCA assay, and serial dilutions were performed with 1% BSA in PBS. Then, 50 μl of the hybridoma supernatant or antibody dilutions were added to individual wells, and the plate was incubated at room temperature for 1 h. The plates were washed with PBS containing 0.1% Tween-20 (PBST_0.1_) three times and then incubated for 1 h with Peroxidase AffiniPure Goat Anti-mouse IgG (H+L) (Jackson ImmunoResearch) (1:5000 dilution). After three washes with PBST_0.1_, TMB signal was generated and detected.

### Pseudovirus neutralization assay

SARS-CoV-2 pseudotyped lentiviruses expressing full-length S proteins and HEK293T cells overexpressing human ACE2 (HEK293T/hACE2) were provided by the National RNAi Core Facility (Academia Sinica, Taiwan). The pseudovirus neutralization assays were performed using serial dilutions of RBD-mAbs pre-incubated with 1000 TU SARS-CoV-2 pseudovirus in 1% FBS DMEM for 1 h at 37°C. The dilutions were incubated for 24 h at 37°C in 96-well white plates (SPL Life Science), which had been pre-seeded with HEK293T/hACE2 cells at a density of 1 × 10^4^ cells per well. The pseudovirus-containing culture medium was then replaced with 10% FBS DMEM for an additional 48 h. Next, ONE-Glo luciferase reagent (Promega) was added to each well for a 3-min incubation at 37°C. Luminescence was measured using a microplate spectrophotometer (Molecular Devices) to determine pseudovirus infection efficacy. The half maximal inhibitory concentration (IC_50_) was calculated by nonlinear regression using Prism software version 8.1.0 (GraphPad Software Inc.).

### Cellular ELISA

HEK293T cells were transiently transfected with wild-type or mutant RBD plasmids in 6-well plates for 16 h. Then, the cells were seeded into 96-well plates at a density of 3 × 10^4^ cells per well and incubated for one day. The cells were fixed in 4% paraformaldehyde in PBS for 15 min at room temperature, and then the cell membranes were permeabilized using 0.1% Triton X-100 at room temperature for 10 min. The plates were blocked using 5% milk, and 100 ng/ml RBD-mAb was added to each well for 1 h at room temperature. Next, the horseradish peroxidase (HRP)-conjugated anti-mouse antibody (1:2000) was added for 1 h at room temperature. The binding capacities of each RBD-mAb to the RBD mutants were examined by ELISA.

### Plaque reduction neutralization test (PRNT)

RBD-mAb-3 was serially diluted in PBS and pre-incubated with 100 plaque-forming units (PFU) of SARS-CoV-2 for 1 h at 37°C. The mixtures were added to pre-seeded Vero E6 cells for 1 h at 37°C. The virus-containing culture medium was then removed and replaced with DMEM containing 2% FBS and 1% methyl-cellulose for an additional 4-day incubation. The cells were fixed with 10% formaldehyde overnight and stained with 0.5% crystal violet for 20 min. The plates were then washed with tap water, and plaque numbers formed were counted for each dilution. Virus without RBD-mAb-3 served as a control. Each experiment was performed in triplicate. Plaque reduction was calculated as: Inhibition percentage = 100 × [1 – (plaque number with mAb / plaque number without mAb)]. The 50% plaque reduction (PRNT_50_) value was calculated with Prism software. The SARS-CoV-2 Delta variant (hCoV-19/Taiwan/ 1144/2021) was obtained from Taiwan Centers for Disease Control (CDC). The PRNT assay was performed in a BSL-3 facility in the Institute of Biomedical Sciences, Academia Sinica.

## FIGURE LEGENDS

**Supplementary Figure 1.**
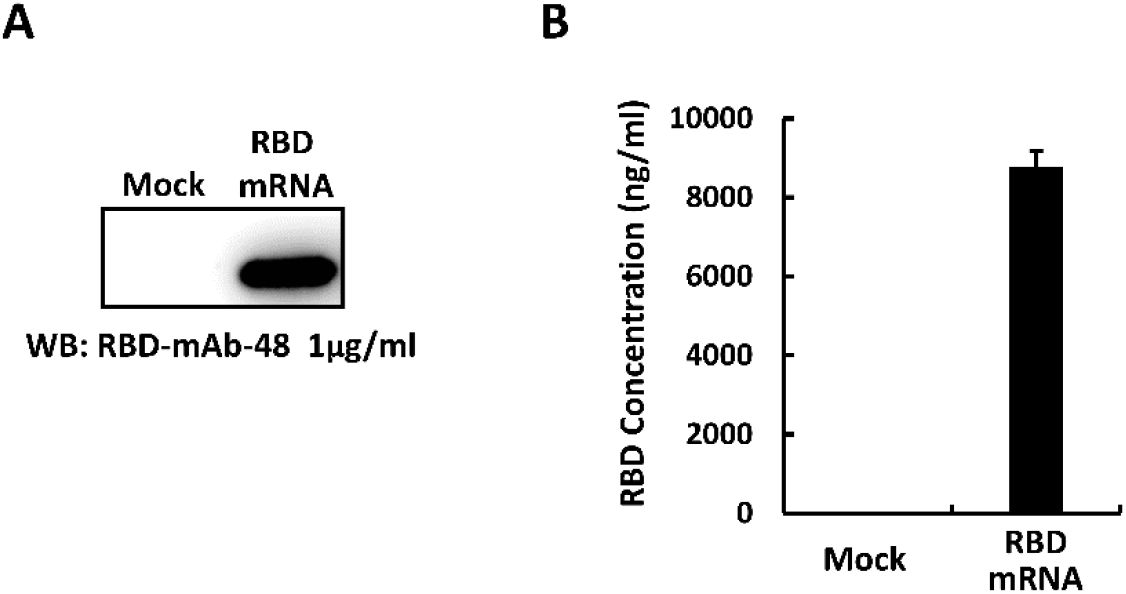
RBD expression from transfected RBD mRNA. RBD protein expression in HEK293T cells transfected with RBD mRNA was measured by Western blotting (A) and ELISA (B), using an anti-RBD antibody; mock transfected HEK293T cells served as a negative control.

**Supplementary Figure 2.**
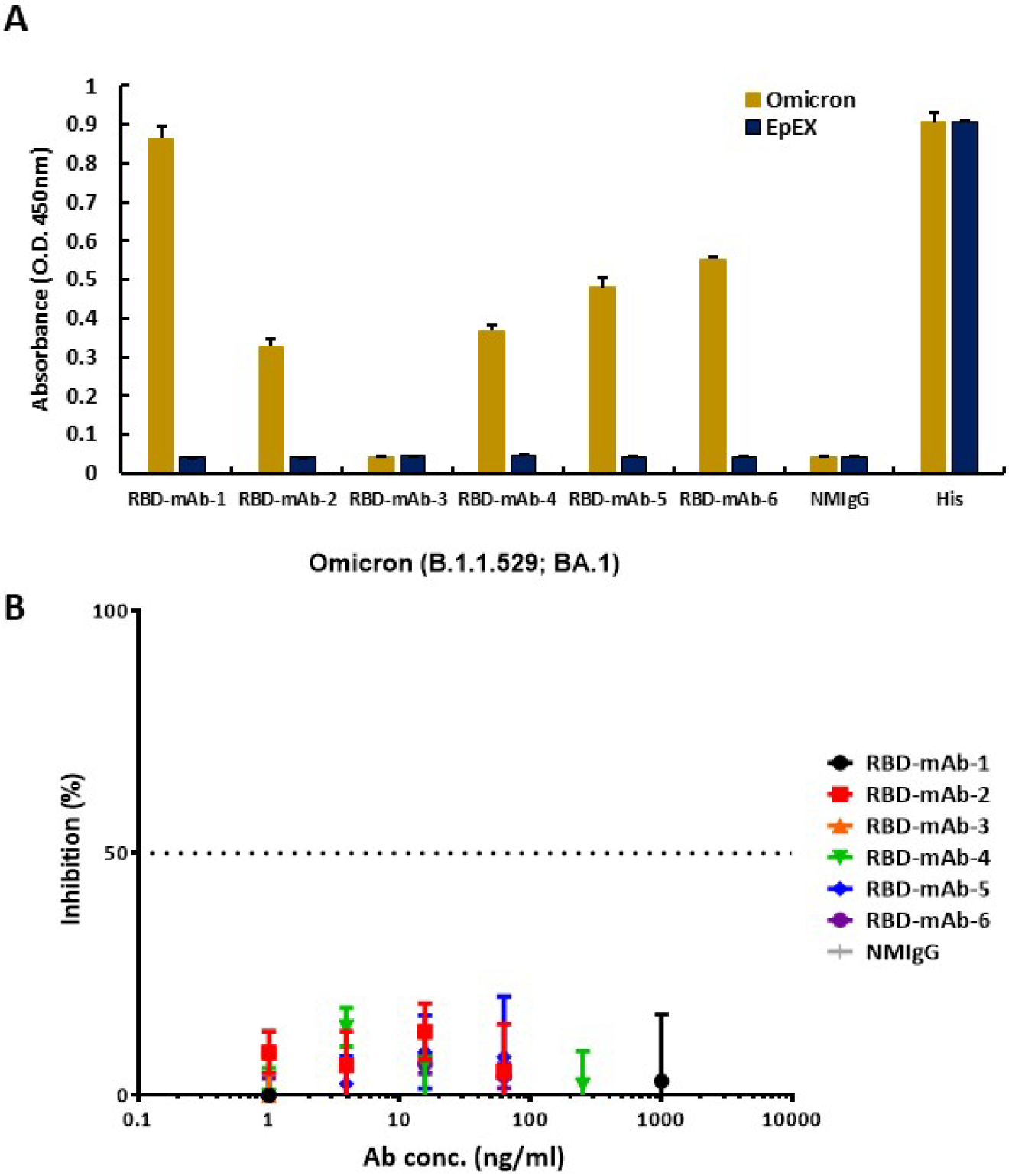
Characterization of Omicron variant binding and neutralizing activities for RBD-mAbs. The binding affinity of hybridoma supernatants to RBD Omicron variant was monitored by ELISA. (B) Neutralization assay of SARS-CoV-2 Omicron pseudovirus with hybridoma culture supernatants. Assays were performed in triplicate, and points represent the mean ± SEM. IC_50_ values were calculated by GraphPad Prism software.

## Acknowledgement

We are indebted to the Human Therapeutic Ab R&D Core facility of BioTReC, Academia Sinica for assisting Ab production and analysis. We also thank for the National RNAi Core Facility at Academia Sinica for providing SARS-CoV-2 variant pseudoviruses.

## Authors’ contributions

FFH designed and performed experiments, and wrote the manuscript. FFH, WYC and HTL immunized the mice and generated mouse mAbs. KHL performed *in vitro* neutralization by pseudoviruses. WYC performed neutralizing Ab expression. FFH, MK, MHT and CMC synthesized mRNA-LNP. HCW conceived the experiments, obtained funding, wrote and revised the paper, and provided overall direction for the study.

## Acknowledgments

We are indebted to the Human Therapeutic Ab R&D Core facility of BioTReC, Academia Sinica, for their assistance with Ab production and analysis. We also thank the National RNAi Core Facility at BioTReC for providing the SARS-CoV-2 variant pseudoviruses.

